# Biotin rescues manganese-induced Parkinson’s disease phenotypes and neurotoxicity

**DOI:** 10.1101/2023.11.21.568033

**Authors:** Yunjia Lai, Pablo Reina-Gonzalez, Gali Maor, Gary W. Miller, Souvarish Sarkar

## Abstract

Occupational exposure to manganese (Mn) induces manganism and has been widely linked as a contributing environmental factor to Parkinson’s disease (PD), featuring dramatic signature overlaps between the two in motor symptoms and clinical hallmarks. However, the molecular mechanism underlying such link remains elusive, and for combating PD, effective mechanism-based therapies are lacking. Here, we developed an adult *Drosophila* model of Mn toxicity to recapitulate key parkinsonian features, spanning behavioral deficits, neuronal loss, and dysfunctions in lysosome and mitochondria. We performed global metabolomics on flies at an early stage of toxicity and identified metabolism of the B vitamin, biotin (vitamin B_7_), as a master pathway underpinning Mn toxicity with systemic, body–brain increases in Mn-treated groups compared to the controls. Using Btnd^RNAi^ mutant flies, we show that biotin depletion exacerbates Mn-induced neurotoxicity, parkinsonism, and mitochondrial dysfunction; while in Mn-exposed wild-type flies, biotin feeding dramatically ameliorates these pathophenotypes. We further show in human induced stem cells (iPSCs)- differentiated midbrain dopaminergic neurons that the supplemented biotin protects against Mn-induced neuronal loss, cytotoxicity, and mitochondrial dysregulation. Finally, human data profiling biotin-related proteins show for PD cases elevated circulating levels of biotin transporters but not of metabolic enzymes compared to healthy controls, suggesting humoral biotin transport as a key event involved in PD. Taken together, our findings identified compensatory biotin pathway as a convergent, systemic driver of Mn toxicity and parkinsonian pathology, providing new basis for devising effective countermeasures against manganism and PD.

**Significance Statement:** Environmental exposure to manganese (Mn) may increase the risk for Parkinson’s disease (PD); however, the mechanistic basis linking the two remains unclear. Our adult fruit fly (*Drosophila*) model of Mn toxicity recapitulated key Parkinson’s hallmarks *in vivo* spanning behavioral deficits, neuronal loss, and mitochondrial dysfunction. Metabolomics identified the biotin (vitamin B_7_) pathway as a key mediator, featuring systemic biotin increases in the flies. Rescue trials leveraging biotin-deficient flies, wild-type flies, and human iPSC-derived dopaminergic neurons determined biotin as a driver of manganism, with the parkinsonian phenotypes dramatically reversed through biotin supplementation. Our findings, in line with overexpressed circulating biotin transporters observed in PD patients, suggest compensatory biotin pathway as a key to untangle the Mn-PD link for combating neurodegenerative disease.

## Introduction

The world is rapidly aging; it is estimated that by 2050, the number of people aged 65 or over will more than double, reaching 1.6 billion which accounts for over 15% of the world’s population (1). These 1.6 billion elderly people will be at substantial risk for brain disease and disorders such as Parkinson’s disease (PD) and Alzheimer’s disease (AD), posing a massive burden on care systems and society due to the long, drawn-out course of disease (1, 2). In the case of treating PD, the most common motor neurodegenerative disease, there are currently no disease-modifying therapies, and for many decades levodopa (L-dopa) has been the only front-line, gold-standard drug since its approval by U.S. FDA in the 1970s (3). The etiology of PD remains elusive and can be diverse; only 10% of PD cases can be explained through Mendelian genetics, suggesting that environmental exposures play a key role in the pathogenesis of the sporadic PD (4).

Manganese (Mn) is an essential micro-nutrient, but chronic exposure to high amounts of it, such as is experienced occupationally by welders, is associated with multiple key motor symptoms of PD. One major pathological hallmark of the disease is aggregation of the protein α-synuclein (αSyn) in neurons—a toxic condition that leads to dysregulated cellular processes and eventually massive neuronal cell death (3). Harischandra et al. found that extracellular vesicles called exosomes isolated from Mn-exposed welders’ blood sera contained misfolded αSyn. In cell culture and rodent models, exposure to Mn or to isolated, Mn-induced exosomes promoted the transfer of αSyn between neurons and microglia, leading to inflammation and eventually to neuronal cell death (5–7). These αSyn-related findings support a causal role of Mn in neurodegenerative disease. Mn has also been linked to induction of neuroinflammation, mitochondrial dysfunction, autophagic defects, and protein aggregation, which all are cellular hallmarks of PD (8). However, there is a dearth of knowledge on the molecular and mechanistic levels regarding how Mn leads to parkinsonism.

Decades of research interrogating individual genes and the molecular basis for PD has led to dozens of ongoing clinical trials aiming to slow down or halt disease progression. However, none thus far have resulted in any effective preventative therapy (9). Here in this study, we took an untargeted approach to identify the early compensatory molecular mechanisms potentially involved in PD pathophysiology. We performed high-coverage, high-resolution mass spectrometry-based metabolomics analyses on an adult fly model of Mn-induced parkinsonism to tease out the early *in vivo* metabolic changes in fly brains and systemically in fly bodies (abdomens and thoraxes) before key parkinsonian phenotypes surfaced. We identified B vitamin biotin (vitamin B_7_) metabolism as a key pathway altered early and showed that supplemented biotin was able to rescue Mn-induced neurotoxicity in both flies and human iPSC-derived dopaminergic neurons including mitochondrial dysfunction. Together, these findings identify compensatory biotin pathway as a systemic, convergent mechanism to resolving the Mn-PD link, providing new basis for developing biotin-based therapies to combat neurodegeneration and environmental health risks.

## Results

### An adult *Drosophila* model of manganese toxicity recapitulates key parkinsonian features

Occupational exposure to manganese has been widely linked as one of the critical environmental factors leading to PD. Although various *Drosophila* models of Mn toxicity have been proposed, most previous models are developmental (10). Since occupational exposure mainly occurs in adults, we developed and characterized an adult model of Mn toxicity in *Drosophila melanogaster*. Flies were fed 1,10- and 30-mM Mn post-eclosion (Fig. 1*A*). Adult exposure to Mn leads to a dose-dependent decrease in lifespan (Fig. 1*B*). In flies post-ten days of Mn exposure, behavioral assays revealed that Mn impaired climbing ability (Fig. 1*C*) and exacerbated locomotor deficits (Fig. 1*D*). In addition, histological analysis at that same time point of the anterior medulla showed that Mn also led to neuronal cell death (Fig. 1*E*). Previous studies by our group and others have shown that Mn induced lysosomal and mitochondrial dysfunction in cell culture and rodent models (6, 7, 11, 12). To validate if the cellular mechanisms of Mn toxicity are consistent in our fly model, we performed the LysoTracker assay on live fly brains. Super-resolution microscopy revealed that Mn led to an increase in lysosomal numbers (Fig. S1A) and size (Fig. S1B), indicating leakage and dysfunction of lysosomes. Further, the Seahorse assay on *Drosophila* brains showed that Mn reduced the oxygen consumption rate (OCR) and drove the cellular metabolic phenotype from energic to quiescent (Fig. 1*G-H*). Finally, the MitoTracker assay demonstrated that Mn led to morphological defects in mitochondria in a dose-dependent manner, as characterized by reduced mitochondrial perimeter and increases in mitochondrial solidity and circularity, which indicated augmented mitochondrial fission (Fig. 1*J-L*). Together, we developed an adult *Drosophila* model of Mn neurotoxicity that recapitulates successfully key parkinsonian features, spanning behavioral deficits, neuronal loss, and dysfunctions in mitochondria and lysosomes.

**Figure 1.**
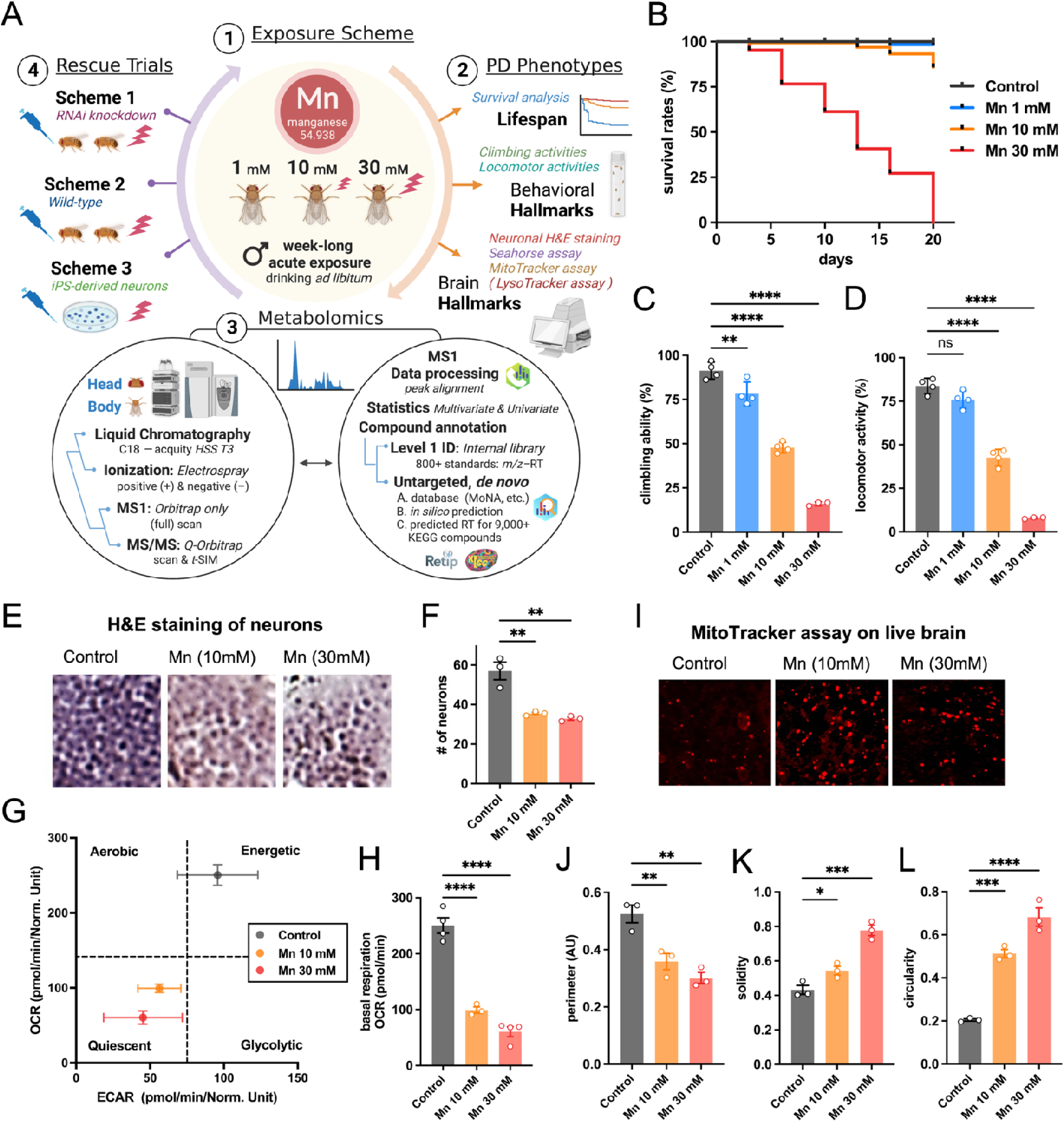
Experimental workflow and the adult *Drosophila* model of manganese (Mn) toxicity that demonstrates behavioral and pathological hallmarks of PD. (*A*) Experimental overview of four modular steps, from exposure scheme, characterization of Parkinson’s phenotypes, metabolomics analysis, to multi-strategy rescue trials. (*B*) Mn exposure reduced the lifespan of *Drosophila melanogaster* in a dose-dependent manner. (*C-D*) Mn exposure led to behavioral deficits in C) climbing and D) locomotor activities in fruit flies. (*E-F*) Histological analysis of anterior medulla by H&E staining revealed that Mn exposure led to significant neuronal loss in brain. (*G*-*H*) The Seahorse metabolic flux assay on live fly brain mapped out (G) a Mn-driven shift in the mitochondrial metabolic phenotype charting the basal respiration oxygen consumption rate (OCR, *x*-axis) and extracellular acidification rate (ECAR, *y*-axis), a measure of lactate formed from glycolytic activities, featuring H) a significant reduction in OCR owing to Mn exposure in a dose-dependent manner. (*I-L*) The MitoTracker assay on live *Drosophila* brain revealed Mn-induced mitochondrial morphological defects as shown by decreased (*J*) mitochondrial perimeter and increases in (*K*) mitochondrial solidity and (*L*) mitochondrial circularity. For statistics, ordinary one-way analysis of variance (ANOVA) was conducted, with data shown as mean ± standard deviation (SD), \**p*<0.05, \*\**p*<0.01, \*\*\**p*<0.001, \*\*\*\**p*<0.0001, Dunnett’s multiple comparisons test. Abbreviations: PD, Parkinson’s disease; iPSCs, induced pluripotent stem cells; HSS, high-strength silica; *t*-SIM, targeted selected ion monitoring; *m/z*, mass-to-charge ratio; RT, retention time; MoNA, MassBank of North America; KEGG, Kyoto Encyclopedia of Genes and Genomes; H&E, hematoxylin and eosin; OCR, oxygen consumption rates.

### High-coverage global metabolomics analysis *in vivo* charts biotin as a key mechanism to Mn toxicity

To identify early biochemical modulators of Mn toxicity, we performed high-coverage untargeted metabolomics analyses of the brain and beheaded body of 5-day-old flies exposed to Mn at low dose (10 mM) and high dose (30 mM) using high-resolution mass spectrometry and integrated cheminformatic computing (Fig. 1*A*; Fig. 2*A*). The rationale for using an earlier time point was to capture the basic biochemical changes that potentially occurred before the behavioral deficits manifested and to identify signature metabolites and/or pathways potentially with a compensatory effect on Mn toxicity. Overall, our untargeted analysis identified distinct metabolome-wide changes owing to Mn exposure, as shown in partial least squares discriminant analysis (PLS-DA) (Fig. 2*B*) and volcano plot (Fig. 2*C*). With high confidence of annotation (level 2 or higher), we resolved 270 and 420 altered metabolites, respectively for brain and systemically for body compartments (Fig. 2*A-B*; *SI Appendix*, Dataset S1-2). We then focused on high dose (30 mM) vs. control only for an in-depth functional analysis, especially of brain metabolome (Fig. 2*A*; Fig. 2*C*), considering that a consistent pattern of changes was seen in body for high dose (30 mM) and low dose (10 mM) relative to control (Fig. S*2*). Our chemical similarity enrichment analysis (ChemRICH) first identified a diverse chemical space of Mn perturbation, enriching altered brain metabolites into 36 chemical classes from moderately nonpolar species such as indoles, succinates, and amino acids towards the more nonpolar terpenes, lysophospholipids, and long-chain fatty acids (Fig. S*3*). Further, quantitative pathway enrichment analysis mapped out a realm of Mn-perturbed pathways, with multiple high-impact ones enriched in B vitamins spanning biotin metabolism, pantothenate and CoA biosynthesis, and vitamin B6 metabolism (Fig. 2*D*). Among the many individual vitamin B family metabolites in fly brain (Fig. 2*E*), biotin embraced the largest fold change (+2.7 fold), the most dramatic statistical significance (lowest *p* value), and a shared trend of change in fly body (+2.0 fold; *SI Appendix*, Dataset S*3*), alongside 50 other metabolites with systemic, brain– body changes (Fig. 2*F*). These together revealed that Mn induced early, systemic metabolic changes, among which biotin metabolism stood out as the top-enriched impact pathway with a compensatory brain–body interaction likely involved.

**Figure 2.**
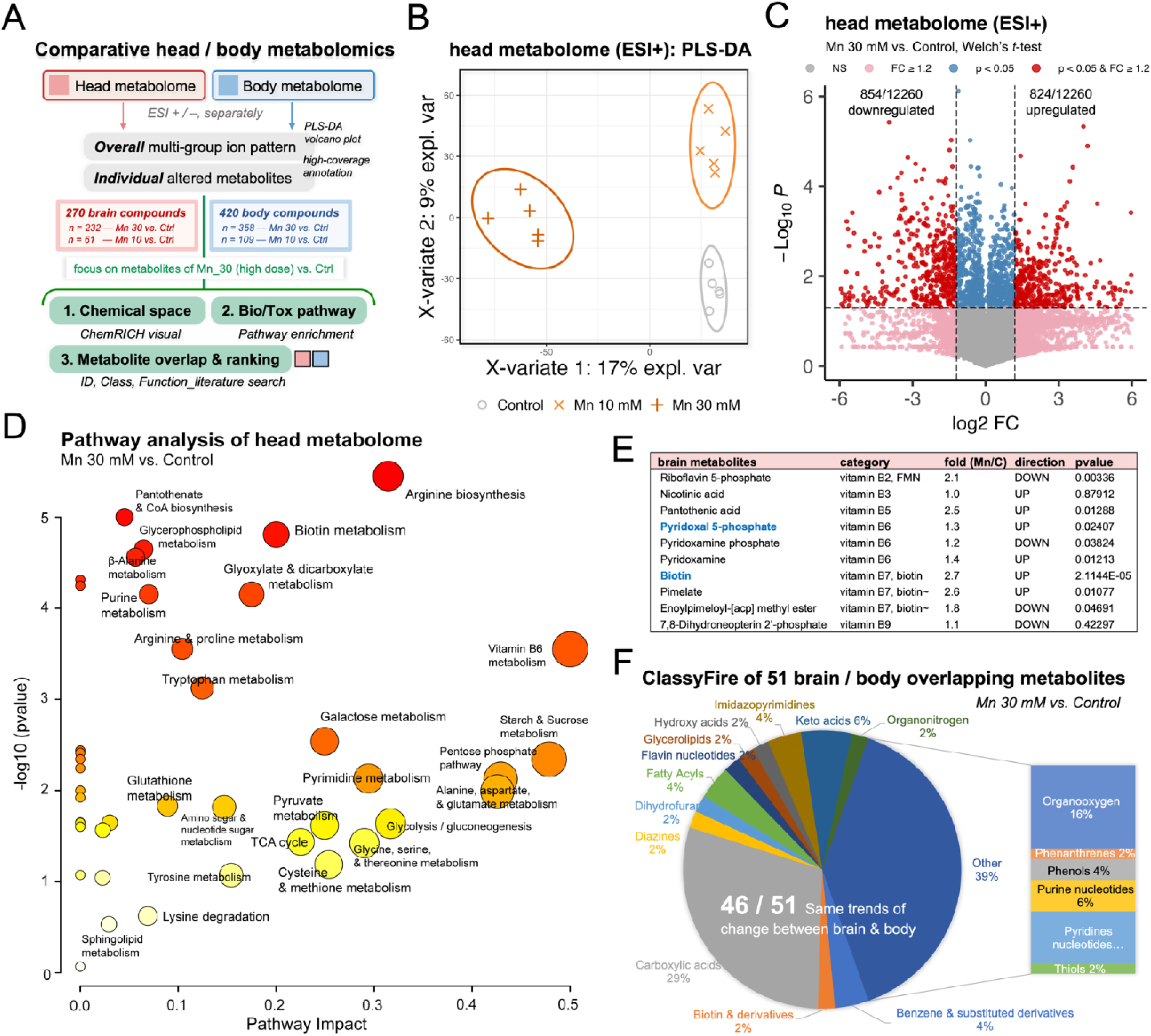
Comparative metabolomics analysis *in vivo* charts biotin metabolism as high-impact and systemic in brain–body interaction in response to Mn exposure. (*A*) Overview of high-coverage comparative metabolomics of head and body compartments of *Drosophila melanogaster* in the Mn exposure scheme, from the overall multivariate pattern to individual metabolites. (*B*) Partial least squares discriminant analysis (PLS-DA) of head metabolomic data for Mn-exposed and control groups acquired under ESI(+) mode; each group has 5 sample replicates with each containing 10 heads. (*C*) Volcano plot of significant ion features as determined by pairwise comparison of high dose (Mn 30 mM) vs. control, Welch’s *t*-test, *p* < 0.05 & fold change ≥ 1.2. (*D*) Pathway analysis of fly head metabolome for high dose (Mn 30 mM) vs. control based on the KEGG *dme* pathway library (for *Drosophila melanogaster*) using the Globaltest approach for pathway enrichment and relative betweenness centrality for the node importance measure in pathway topological analysis. (*E*) Multiple vitamin B family metabolites altered in *Drosophila* head metabolome comparing high dose (Mn 30 mM) vs. control, with biotin embracing the most dramatic statistical significance (lowest *p* value), largest fold change (+2.7 fold), and a shared direction of change in fly body (+2.0 fold, not shown in chart); blue bold highlights metabolites altered in both fly brain and body. (*F*) ClassyFire pie chart view of 51 brain–body overlapping metabolites comparing high dose (Mn 30 mM) vs. control, Welch’s *t*-test, *p* < 0.05 & fold change ≥ 1.2; 46 of the total 51 shared the same directions of change between brain and body, including biotin. Abbreviations: ESI, electrospray ionization; ChemRICH, chemical similarity enrichment analysis; FC, fold change; FMN, flavin mononucleotide.

### Biotin drives Mn-induced neurotoxicity, parkinsonism, and mitochondrial dysfunction

In light of our metabolomics findings of biotin, we undertook experimental studies to test systematically for its functional roles in Mn-related neurodegeneration. Biotin is an essential water-soluble B_7_ vitamin and can either be recycled, obtained from diet, or produced by the gut microbiome (13). The biotinylation of carboxylases has been widely implicated in cellular functions including mitochondrial respiration (14) and neurotransmitter production (15, 16), among others. Most importantly, biotin is essential for dopamine production (16). Therefore, we hypothesized that Mn triggers off biotin recycling as a compensatory mechanism at an early stage before parkinsonism manifests. To test this hypothesis, we knocked down *Btnd*, the gene responsible for biotin recycling, specifically in fly neurons using RNA interference (RNAi), and then exposed the mutant flies to Mn (Fig. 3*A*). Climbing (Fig. 3*B*) and locomotor assay (Fig. 3*C*) both showed that neuronal *Btnd* knockdown further exacerbated Mn-induced behavioral deficits compared to Mn-exposed wildtypes. Moreover, the MitoSox assay on live fly brains determined that diminished levels of free biotin boosted mitochondrial superoxide production in Mn-exposed flies (Fig. 3*D*). Finally, the Seahorse metabolic assay on *Drosophila* brains determined that the neuronal knockdown of *Btnd* lowered mitochondrial oxygen consumption rates (Fig. 3*E*) and the overall cellular phenotype (Fig. 3*F*) in flies fed with Mn.

**Figure 3.**
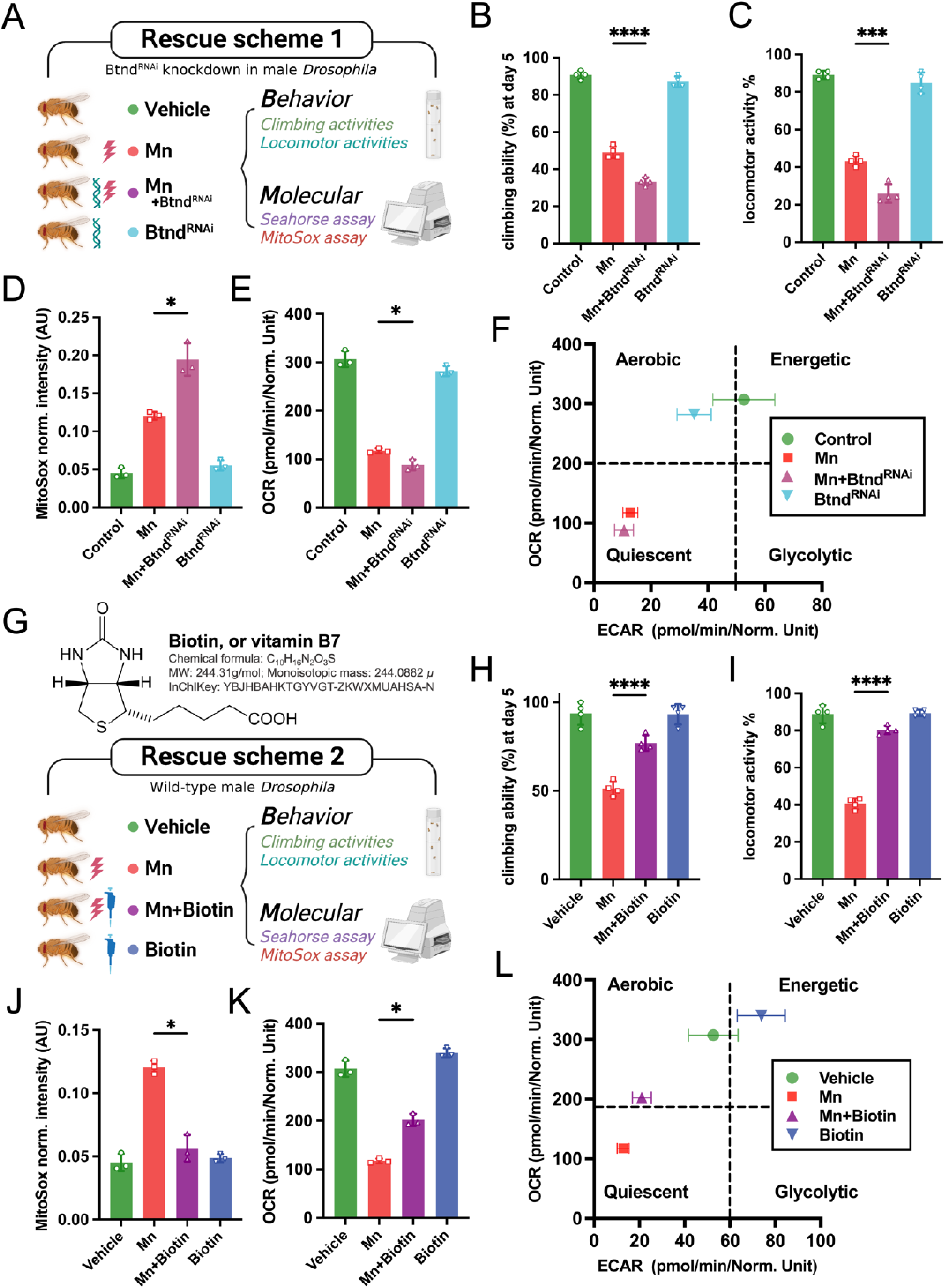
Rescue trials *in vivo* demonstrate biotin as an essential agent that protects against Mn-induced damage and parkinsonism. (*A*) Schematic of Rescue Trial 1 *in vivo* including Btnd^RNAi^ mutant male flies (knockdown of neuronal *Btnd* gene encoding biotinidase) exposed to Mn (30 mM). (*B-C*) Biotin deficiency exacerbated Mn-led behavioral deficits in B) climbing and C) locomotor activities in fruit flies. (*D*) The MitoSox assay on live *Drosophila* brain revealed that biotin deficiency dramatically heightens Mn-induced superoxide levels. (*E-F*) The Seahorse metabolic flux assay on live fly brain determined for Btnd^RNAi^ mutant lines that biotin deficiency exacerbated E) Mn-induced reduction in mitochondrial OCR and led to F) a more dysfunctional shift of metabolic phenotype under Mn exposure (30 mM) compared to wild-type files. (*G*) Schematic of Rescue Trial 2 *in vivo* using wild-type male flies exposed to Mn (30 mM) with biotin supplementation. (*H-I*) Biotin feeding significantly recovered Mn-led behavioral deficits in H) climbing and I) locomotor activities in wild-type fruit flies. (*J*) The MitoSox assay on live *Drosophila* brain revealed that biotin feeding significantly reduced mitochondrial superoxide levels caused by Mn exposure. (*K-L*) The Seahorse metabolic flux assay on live fly brain determined that supplemented biotin significantly recovered K) the mitochondrial OCR and L) ameliorated the shifted damages of metabolic phenotype induced by Mn exposure. For statistics, ordinary one-way analysis of variance (ANOVA) was conducted, with data shown as mean ± standard deviation (SD), \**p*<0.05, \*\**p*<0.01, \*\*\**p*<0.001, \*\*\*\**p*<0.0001, Dunnett’s multiple comparisons test. Abbreviations: RNAi, RNA interference; good. HMDB, human metabolome database.

To investigate whether the compensating biotin has protective or preventive potential, we supplemented biotin into fly diet through which the adult flies were co-exposed to Mn (Fig. 3*G*). Both the climbing (Fig. 3*H*) and locomotor assays (Fig. 3*I*) demonstrated that biotin feeding ameliorated Mn-induced behavioral deficits. The MitoSox assay on *Drosophila* brains showed that dietary biotin supplementation inhibited mitochondrial superoxide production in Mn-exposed flies, virtually lowering to the levels of non-Mn groups (Fig. 3*J*). Functionally, the Seahorse assay on live fly brains revealed that biotin rescued mitochondrial oxygen consumption rates (Fig. 3*K*) and the cellular phenotype (Fig. 3*L*) in Mn-exposed flies. In parallel, histological analysis determined that while a reduction in free biotin pool through *Btnd* knockdown exacerbated Mn-induced neuronal loss, biotin supplementation protected against it (Fig. S*4*). In a nutshell, these data suggested that free, bioavailable biotin plays a crucial compensatory role in regulating neurodegeneration induced by Mn *in vivo*.

### Biotin ameliorates Mn-induced neurotoxicity in iPSC-derived midbrain dopaminergic neurons

Biotin deficiency has been implicated in mitochondrial dysfunction and neuronal loss (14). To validate our findings with mammalian relevance for its therapeutic potential, we differentiated human induced pluripotent stem cells (iPSCs) into midbrain dopaminergic neurons and aged them for 20 days. Post-aging, we pre-treated these cells with biotin for 24 hours and then exposed them to Mn for 24 additional hours. Super-resolution microscopy detected that biotin supplementation protected profoundly against Mn-induced tyrosine hydroxylase (TH) neurite loss (Fig. 4*A-B*). Both the LDH assay (Fig. 4*C*) and MTS assay (Fig. 4*D*) determined that biotin inhibited Mn-induced cytotoxicity in these iPSC-derived dopaminergic neurons. Moreover, the MitoTracker green assay and MitoSox assay revealed respectively for these neuronal cells that biotin rescued Mn-led mitochondrial mass loss indicative of impaired mitochondrial function and biogenesis (Fig. 4*E*) while lowering mitochondrial superoxide production (Fig. 4*F*). Finally, to gauge cellular functional protection, we performed the Seahorse assay on biotin-pretreated dopaminergic neurons after Mn exposure (Fig. 4*G-H*). Supplemented biotin markedly restored Mn-induced loss of oxygen consumption rate (OCR) in dopaminergic neurons (Fig. 4*G*) and rectified partially the quiescent, glycolytic phenotype caused by Mn exposure (Fig. 4*H*). Meanwhile, to illuminate the role of biotin in human PD, we conducted data mining in NCBI Gene Expression Omnibus (GEO) for biotin-related genes and their expressions. We discovered that compared to age-matched healthy controls, PD patients had an elevated circulating level of biotin transporters but not biotin-metabolizing enzymes in the brain (Fig. 4*I*), suggesting for PD an altered pattern in biotin transport. Our results together showed that biotin supplementation protected human iPSC-derived dopaminergic neurons from Mn-induced neurotoxicity and mitochondrial dysfunction, with the PD patient data indicating elevated biotin transport as a key event in the pathogenesis of PD.

**Figure 4.**
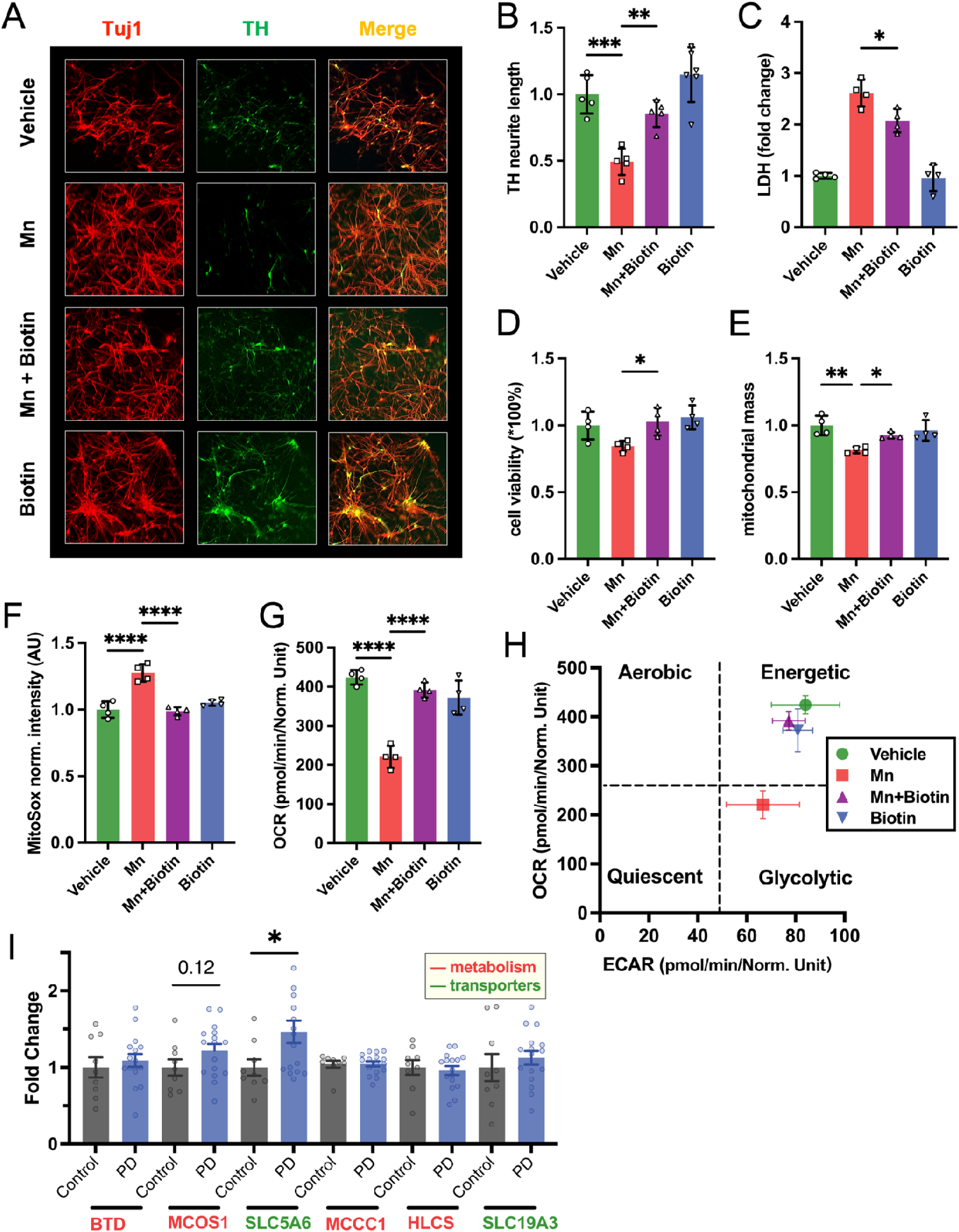
Biotin as a convergent mechanistic player linking manganism to Parkinson’s disease: experimental rescue trials using human iPSC-derived midbrain dopaminergic neurons and relevant protein expression levels reported for PD patients. (*A-C*) Rescue trials *in vitro* using iPSC-derived midbrain dopaminergic neuronal cell culture demonstrated that supplemented biotin protected against dopaminergic neuronal loss induced by Mn, as shown for the biotin-fed group in A) tyrosine hydroxylase (TH) staining (middle green panel), a marker for dopaminergic neurons, B) recovered TH neurite length from Mn exposures, and C) decreased LDH levels, an indicator of neuronal membrane damage caused by Mn exposure. (*D*) The MTS assay determined that biotin supplementation recovered Mn-reduced cell viability for iPSC-derived neurons. (*E*) The MitoTracker assay on live *Drosophila* brain determined that biotin supplementation significantly restored the Mn-induced mitochondrial mass loss in dopaminergic neurons. (*F*) The MitoSox assay revealed that biotin feeding reduced mitochondrial superoxide levels induced by Mn in iPS-derived neurons. (*G-H*) The Seahorse metabolic flux assay on iPSC-derived neurons maps for biotin-fed groups G) a dramatic recovery from Mn-induced metabolic phenotypic shift and that H) biotin significantly ameliorated Mn-inhibited neuronal mitochondrial OCR. (*I*) Human PD patient data on biotin-related protein expression levels (fold change are normalized values of all) showed for PD cases vs. control differentiated levels in biotin transporters (e.g., SLC5A6 and SLC19A3) but not in biotin metabolic enzymes; all data were retrieved from NBCI Gene Expression Omnibus (GEO) database including *Homo sapiens* data only (last accessed in April 2023). For iPSC cell data, ordinary one-way ANOVA with Dunnett’s multiple comparisons test was conducted, with data shown as mean ± standard deviation (SD); for human gene expression data, Student’s *t*- test was conducted for each protein comparing PD case and control, with data shown as mean ± standard deviation of the mean (SEM), \**p*<0.05, \*\**p*<0.01, \*\*\**p*<0.001, \*\*\*\**p*<0.0001. Abbreviations: iPSC, induced pluripotent stem cells; Tuj1, neuron-specific class III β-tubulin; LDH, lactate dehydrogenase; BTD, biotinidase; MCOS1, molybdenum cofactor synthesis 1; SLC5A6, sodium-dependent multivitamin transporter, or solute carrier family 5 member 6; MCCC1, methylcrotonyl-CoA carboxylase subunit 1; HLCS, holocarboxylase synthetase; SLC19A3, thiamine transporter 2, or solute carrier family 19 member 3.

## Discussion

Among the realm of brain diseases that increasingly plague the aging world, Parkinson’s disease is the most common motor disorder (2). Since its first medical documentation in 1817, PD has long attracted research interests as a pathfinder disease to gather clues for resolving idiopathic neurodegeneration altogether. However, disease-modifying therapies are lacking, and the molecular mechanism of PD still remains elusive with the environmental risk contributors largely unexplored (4). To identify and develop preventative strategies for PD, in this study, we performed high-coverage, untargeted metabolomics on an adult *Drosophila* model of manganese toxicity at an early timepoint over parkinsonian phenotypic manifestation. We identified biotin metabolism as a top-enriched pathway featuring systemic increases of biotin in both the brain and body of flies exposed to Mn. Using genetics and pharmacological approaches, we demonstrated *in vivo* and *in vitro* that biotin can alter the course of Mn-induced neurotoxicity through the mediation of mitochondrial dysfunction.

Exposure to occupational, neurotoxic metals like Mn has been linked to the development of PD (8). Previous studies have demonstrated that Mn can bind to the metal binding sites of α-synuclein (αSyn) and lead to aggregation of this protein, one of the key pathological hallmarks of PD. At an earlier time, Mn may protect the neurons against αSyn toxicity, but overexposure to Mn likely enhances the neuronal accumulation and propagation of αSyn (5, 6). Further, Mn can also intensify αSyn-induced inflammation, another hallmark of PD (7, 17). Mn has also been recently shown to induce the expression of leucine-rich repeat kinase 2 (*LRRK2*), another key *PARK* gene that has been linked to PD etiology (18, 19). For disease modeling, previous fly models of Mn toxicity were largely based on the developmental larvae, lacking relevance for occupational exposures to Mn occurring in adults, from whom the Mn-PD links were first drawn (10). Here, we thus developed an adult *Drosophila* model of Mn neurotoxicity that recapitulates various key aspects of PD, including mitochondrial dysfunction, lysosomal defects, behavioral deficits, and neuronal loss.

PD is a multi-system disorder, as it concomitantly involves many non-motor symptoms that arise even decades before PD’s cardinal motor symptoms surface (20); one of the earliest non-motor symptoms is constipation. Braak’s hypothesis of PD states that sporadic PD may initiate in the gut (21). In PD patients, the αSyn pathology has been detected in the gut at the early stages of the disease (22, 23). Further, studies in rodent models have demonstrated that αSyn originating from the gut (converted *in situ* from αSyn fibrils injected in) can spread to the brain through the vagus nerve in a prion-like manner, leading to features of PD (24). Mn is known to cause autonomic dysfunctions similar to PD (11), and the recent studies by Payami and coworkers have associated PD with widespread microbiome dysbiosis (25, 26). In this light, to better understand Mn toxicity and its links to PD, we took a non-targeted, multi-compartmental approach to probe metabolome-wide changes *in vivo* at an early timepoint of Mn exposure before the motor symptom sets in. Using high-resolution mass spectrometry, we conducted high-coverage, untargeted metabolomics to identify molecular signatures in brain and systemically in the body of flies owing to Mn exposure. Strikingly, annotated with confidence, more than 400 metabolites were altered in the body, and over 250 metabolites were altered in the brain. Given our hypothesis considering PD as a multi-system disorder, we compared these metabolite profiles across datasets and identified that, among the multiple B vitamin pathways significantly perturbed in the brain, biotin metabolism stood out as the most high-impact pathway with systemic, body–brain increases in Mn-treated flies.

Biotin, or vitamin B_7_, is a water-soluble essential vitamin either obtained through diet or produced by the gut microbiome (13). Biotin acts as a critical cofactor for various carboxylases involved in a range of metabolic processes, including dopamine production (27). Previous studies have implicated that biotin deficiency can cause locomotor deficits and mitochondrial pathology in a tau-toxicity model (14). Such biotin deficiency in rats also leads to short-term memory loss and deteriorated locomotor activity, further suggesting a role of B-vitamin biotin in PD (27). Gut microbiota is known to mediate B vitamin bioavailability; a recent study showed that prebiotic supplementation increases the abundance of biotin-producing gut bacteria, particularly *Bacteroides* spp., in a high-fat diet mouse model (13). Interestingly, of the *Bacteroides* genus, many biotin-related genes and one specific species were also altered in human PD patients’ fecal samples (25). Furthermore, a recent metabolomic study in the A53Tα-synuclein mouse model of PD identified biotin metabolism as a key altered pathway (28). In this study, we show that biotin deficiency exacerbates Mn-induced neurotoxicity in both flies and in patient-originated, iPSC-derived midbrain neuronal cell cultures. By contrast, biotin supplementation dramatically inhibits all these pathophenotypes induced by Mn exposure.

Mitochondrial dysfunction has been recognized as a key molecular mechanism of pathogenesis in various PD models (29–33). Research has shown that the hallmark α-Syn protein can bind to mitochondria and alter their cellular function directly (34). Further, α-Syn also induces actin hyperstabilization, which can affect the mitochondrial fission-fusion process (35). In parallel, mutation of other *PARK* genes, such as *LRRK2*, has been demonstrated to induce mitochondrial dysfunction in animal and cellular models as well (36). Here, we further show *in vivo* and *in vitro* that biotin supplementation is able to rectify Mn-induced neurotoxicity through rescue of mitochondrial dysfunction in neurons, spanning aspects of mitochondrial mass, morphology, respiration, and the overall metabolic phenotypes. Future research is thus warranted to further decipher the molecular underpinnings (e.g., biotin transport, ligand binding sites) of biotin-mitochondria interaction for PD and many other pertinent neurodegenerative conditions.

The therapeutic potential of B-vitamin biotin as a countermeasure against PD and neurodegeneration is immense. Biotin is well tolerated in humans, and with new microbiota manipulation technologies in place (37), novel, non-pharmacological approaches such as supplementation through biotin-rich prebiotics and/or biotin-producing probiotics may be an effective avenue for curbing PD. Given that autonomic dysfunction is an early key symptom widely involved in PD pathogenesis and aging populations in general, future studies are needed to dissect the role of gut dysbiosis in regulating biotin production and recycling, risks of environmental neurotoxic exposures, and the neurodegenerative conditions altogether.

To summarize, we applied a high-coverage, untargeted metabolomics using high-resolution mass spectrometry and advanced cheminformatics computing in a newly developed adult model of metal-induced parkinsonism in *Drosophila melanogaster* at the forefront of PD research. This non-targeted approach generates an unbiased, comprehensive metabolomics dataset at the metabolite levels that is of broad interest, leading us to the discovery of biotin metabolism as an early master compensatory modifier of Mn-induced neurodegeneration. The findings and omics datasets of this study may serve as a powerful resource and example for the identification of actionable therapeutic targets from systems-level analysis of organismal disease models.

## Materials and Methods

### *Drosophila* husbandry and feeding

Fly crosses were conducted in a 25 °C incubator and aged at 25 °C for 5-20 days, depending on the experiments. The pan-neuronal driver nSyb-GAL4 was used to mediate the genetic knockdown of the *Btnd* gene. Both the stocks for nSyb-GAL4 and UAS-Btnd RNAi were obtained from Bloomington Drosophila Stock Center (BDSC) (HMC05012; Bloomington #60020). For Mn treatment, 0, 1, 10 and 30 mM of MnCl_2_ (#7773-01-5; Santa Cruz Biotechnology, Dallas, TX) was mixed in 3 mL of water, and the instant food for flies was made using this water. Biotin supplementation was performed as previously described (14). Biotin was added to instant fly food (Carolina Biological, Burlington, NC) at a final concentration of 30 mM. Unless noted otherwise, male flies were used throughout the study.

### iPSC cell culture and treatment

Human midbrain dopaminergic neurons were differentiated from induced pluripotent stem cells (iPSCs) purchased from Axol Biosciences (Cambridge, UK). The cells were differentiated and matured using the StemCell Midbrain Neuron differentiation and maturation kit (#100-0038 and #100-0041; StemCell Technologies, New York, NY). Briefly, 24- and 96-well plates were coated with laminin and poly-L-ornithine. Cells were then plated into StemCell Stemdiff^TM^ Midbrain Neural Differentiation Media (Mid-NDM), which was supplemented with 200 ng/ml human recombinant-Shh and 10-μM ROCK inhibitor. Every other day for seven days, half of the media was refreshed. Post seven days, we switched to using BrainPhys^TM^ Neuron Maturation Medium (#100-0041; StemCell Technologies, New York, NY), with half of the media being replaced every alternate day. All treatments were diluted using the BrainPhys^TM^ Neuron Maturation Medium; for treatment groups, 30 μM biotin and 100 μM Mn were supplemented in the BrainPhys Neuron Maturation Medium for 24 hrs.

### Climbing and locomotor assay

The climbing and locomotor activities were assessed following previously published procedures (36, 38, 39). To gauge climbing ability, we counted the number of flies climbing 5 cm within 10 secs. In each vial, at least ten flies were present, and for each group, we had four vials. For the locomotor assay, each vial containing over 10 flies was tapped and placed horizontally. After 15 seconds, the number of flies walking was counted (40). For two groups, data were analyzed using Student’s *t*-test; for multiple groups, one-way ANOVA with *post hoc* analysis was performed.

### Histological assessment of neuronal loss

Fly brains were fixed with formalin and paraffin embedded. Then, 2 μm sections were cut, and hematoxylin and eosin (H&E) staining was performed, imaged using the Nikon EclipseE600 (Melville, NY), and counted using ImageJ as previously described (36, 41).

### LysoTracker assay

The LysoTracker assay (kit #L7526; Thermo Fisher Scientific, Waltham, MA) was conducted as described (39) to assess lysosomal morphology. Briefly, fresh whole-mount brains from 10-day-old flies were incubated with 1 μM LysoTracker for 5 min at room temperature, mounted in PBS, and imaged immediately on a Zeiss laser-scanning confocal microscopy (Oberkochen, Germany). The number and size of the lysosomes were analyzed using ImageJ.

### MitoTracker assay

Fly brains were dissected and incubated in 1-μM MitroTracker red dye (kit #M-7512, Thermo Fisher Scientific, Waltham, MA) for 15 mins, washed in PBS twice, and imaged using a Zeiss laser-scanning confocal microscopy (Oberkochen, Germany). The solidity, perimeter, and circularity were measured following previously published protocols (32). For iPSCs, mitochondrial mass was measured using a published protocol (17). Briefly, post-treatment, cells were incubated in 1-μM MitroTracker green dye (kit #M7514, Thermo Fisher Scientific, Waltham, MA) for 13 mins, washed using HBSS, and placed on a plate reader to quantify the fluorescence. Then, the values were normalized using nuclear stain, DAPI, and fluorescence.

### Seahorse assay

Brain metabolic changes in *Drosophila* were measured using the Seahorse XFe96 metabolic bioanalyzer (North Billerica, MA). Oxygen consumption rates (OCRs) and extracellular acidification rates (ECARs) were determined as described previously (38). For all experiments, brains from 5-day-old flies of the appropriate genotypes were dissected and plated at one brain per well on XFe96 plates (Seahorse Bioscience, North Billerica, MA), and metabolic parameters were assayed as described (36, 38). The OCR values were normalized to DNA content using a CyQUANT assay (Thermo Fisher Scientific, Waltham, MA) following the manufacturer’s instruction. For cells, the Seahorse assay was performed using our previous paper (3) for cells. Post-treatment, the Seahorse plate was incubated in a CO_2_-free incubator for an hour. The calibration plate was incubated overnight in the CO_2_-free incubator at 37°C. The ECAR and OCAR values were normalized using CyQUANT (kit #C7026, Thermo Fisher Scientific, Waltham, MA).

### MitoSOX assay

Mitochondrial reactive oxygen species (ROS) activity was assessed using the MitoSOX kit (#M36008, Thermo Fisher Scientific, Waltham, MA) for fly brains and iPSC-derived TH neurons (32, 36). Fly brains and iPSC cells were incubated in 1-μM MitoSox red dye (kit #M7514, Thermo Fisher Scientific, Waltham, MA) for 15 mins, washed using PBS, and a plate reader was used to quantify the fluorescence. Then, the values were normalized using nuclear stain, DAPI, and fluorescence.

### Immunocytochemistry (ICC)

ICC was performed using previously described protocols (33, 38, 42). Briefly, cells were plated and treated on a coverslip in 24-well cell culture plates. Post-treatment, cells were gently washed and fixed using 4% paraformaldehyde for 30 mins, washed with PBS twice, blocked in blocking buffer (2% BSA, 0.5% Triton-X, 0.05% Tween), and incubated with primary antibodies overnight at 4°C. Post-incubation, cells were washed with PBS, incubated with secondary antibodies, washed again, and mounted on slides using a DAPI-containing mounting medium. The primary antibodies used in this study included Anti-Tyrosine Hydroxylase Antibody (#AB152, RRID:AB_390204; Millipore Sigma, Burlington, MA) and Purified anti-Tubulin β 3 (TUBB3) Antibody (#801201, RRID:AB_2313773; BioLegend, San Diego, CA).

### LDH cytotoxicity assay

A total of 100 μL of conditioned media was collected from cultures treated with manganese. The lactate dehydrogenase (LDH) release was measured using LDH Assay Kit (Fluorometric) (#ab197000; Abcam, Cambridge, UK) following the manufacturer’s protocol.

### MTS cell viability assay

Cells were plated in 96-well tissue plates. After treatment, 10 µL of 3-(4,5-dimethylthiazol-2-yl)-5-(3-carboxymethoxyphenyl)-2-(4-sulfophenyl)-2H-tetrazolium (MTS) reagent (#G3582; Promega, Madison, WI) was added and incubated at 37°C for 45 mins. After incubation, a plate reader was used to quantify the absorbance at 490 nm. Then, the values were normalized using nuclear stain, DAPI, and fluorescence (17).

### Metabolomics analysis

To capture basic biochemical changes, flies were beheaded at 5-day of Mn exposure with heads and bodies collected and snap-frozen separately; each group had five sample replicates, and each replicate sample contained ten heads/bodies. To extract metabolites, thawed heads and bodies were added with ice-cold methanol:H_2_O (2:1, *v/v*) solution pre-spiked with isotope-labeled internal standards alongside zirconium oxide beads (0.5 mm i.d., Yittria stabilized) (Next Advance, Troy, NY). Samples were touch-vortexed for 30 sec, manually ground using sterile, disposable pestles (DWK Life Sciences, Millville, NJ), placed on a bead beater (Next Advance, Troy, NY) in a cold room for 5 mins at the maximum speed, and finally centrifuged at 18,000 × *g* for 10 mins. The supernatant was SpeedVac-dried and reconstituted into acetonitrile:H_2_O (2:98, *v/v*) upon instrumental analysis. As detailed elsewhere, a 15-min method of C18 liquid chromatography was used (43), followed by fullscan high-resolution MS1 data acquisition using a Thermo Vanquish UHPLC coupled to an Orbitrap Exploris^TM^ 240 high-resolution mass spectrometer interfaced with a heated electrospray ionization (ESI) source (Waltham, MA). Data in both ESI positive and negative modes were acquired. Quality assurance and quality control (*QA/QC*) procedures were implemented, spanning timely mass calibration, sample blindfolding, sample randomization, and internal standard-based monitoring. MS1 *.raw data were converted to *.abf, and processed in MS-DIAL 4.90 (Riken, Japan) (44) to obtain peak alignment tables separately for head (ESI+), head (ESI-), body (ESI+), and body (ESI-) modes of analysis; detailed settings for data processing can be found in the *SI Appendix*. Welch’s *t*-test and one-way ANOVA were performed on each of the four datasets to screen for ion features of statistical significance; retrospectively, tandem MS/MS mass spectra were collected for these features correspondingly using pooled samples, separately for fly heads and bodies.

### Informatics

To identify chemical structures, a three-pronged cheminformatic approach was applied to reach the highest compound coverage possible, featuring an integrated strategy of (i) matching against an internal RT-*m/z* library of 812 common metabolites established from authentic chemical standards (IROA Technologies, Bolton, MA), (ii) *de novo* formula and structural prediction in MS-FINDER 3.30 (Riken, Japan) (45) based on hydrogen rearrangement rules, using accurate mass, isotope ratios, and tandem MS/MS of ions of interest as input data, and (iii) matching against an *in silico* RT-*m/z* library constructed through pairing theoretical accurate mass (of [M+H]+ for ESI+ and of [M-H]- for ESI-) with machine learning-based predicted chromatographic retention time for over 9,000 relevant structures compiled from KEGG, a recently reported atlas of mouse brain metabolites (46), and a newly curated list of microbiome metabolites from Exposome Explorer (47). The confidence of annotation was assigned to each structure based on the Schymanski Scheme in compliance with the Metabolomics Standard Initiative (MSI) guidelines (48). To infer metabolomic patterns and pathways, multivariate statistics, chemical enrichment analysis, and quantitative pathway analysis were performed. Multivariate partial least squares discriminant analysis (PLS-DA) was conducted in R (Vienna, Austria) using the *mixOmics* R package. Chemical similarity enrichment analysis (ChemRICH) plots were constructed on altered metabolites to visualize the chemical space as clustered/enriched by compound classes (Davis, CA). Pathway enrichment analysis was performed in MetaboAnalyst 5.0 (Quebec, Canada) based on the KEGG *dme* pathway library (organism-specific, *Drosophila melanogaster*), using the Globaltest approach for pathway enrichment and relative betweenness centrality for the node importance measure in pathway topological analysis (49).

### Human gene expression data

Datasets of human PD were retrieved from the NCBI GEO database with the MeSH search terms “parkinson disease” OR “Parkinson’s diseases” AND “gds” (for DataSets) AND “Homo sapiens,” resulting in 12 human studies of PD in total with deposited datasets of gene expression profiling of blood samples by array to be included in our pooled, pairwise comparison of PD vs. healthy controls. We examined the circulating expression levels for two types of biotin-related genes, including key enzymes for biotin metabolism (e.g., BTD, MCOS1, MCCC1, HLCS) and biotin transporters (e.g., SLC5A6, SLC19A3). Fold change was calculated for PD levels as normalized to controls.

### Statistical analysis

Unless otherwise stated, all sample sizes (*n*) indicate biological replicates rather than technical replicates. The adopted sample sizes were determined based on our previous work handling these models and techniques. For all behavioral and molecular phenotyping data, we used ordinary one-way ANOVA with Dunnett’s multiple comparisons test for multiple comparisons and unpaired Student’s *t*-test for comparison between two groups in GraphPad Prism 9 (San Diego, CA). For metabolomics data, Welch’s *t*- test was conducted for pairwise comparison, and one-way ANOVA and Tukey’s HSD test were used for *post hoc* multigroup comparison.

## Acknowledgments

This work was funded by the National Institutes of Health (NIH) through awards to G.W.M. (R01ES023839, U2C ES030163, UL1 TR001873, R01AG067501, and RF1AG066107) and to S.S. (R00 ES033723 and P30 ES001247). The authors thank the instrumentation and dedicated staff support from the Exposomics Laboratory and the Irving Institute Biomarkers Core Laboratory (BCL) at the Columbia University Irving Medical Center.

## Data sharing plans

The metabolomics raw data will be deposited in the National Metabolomics Data Repository (NMDR) through Metabolomics Workbench. R codes, if necessary, can be made publicly available in GitHub and Zenodo. Source data for figures in this study can be made available upon request by the editorial board, reviewers, or prospective readers.

## Notes

### Competing Interest Statement

The authors have declared no competing interest.

## References

1. E. United Nations. Department of, A. Social, World Social Report 2023 : Leaving No One Behind in an Ageing World (United Nations, New York, 2023).

2. G. U. N. D. Collaborators et al., Burden of Neurological Disorders Across the US From 1990-2017: A Global Burden of Disease Study. JAMA Neurol 78, 165–176 (2021).

3. M. J. Armstrong, M. S. Okun, Diagnosis and Treatment of Parkinson Disease: A Review. JAMA 323, 548–560 (2020).

4. A. Ascherio, M. A. Schwarzschild, The epidemiology of Parkinson’s disease: risk factors and prevention. Lancet Neurol 15, 1257–1272 (2016).

5. D. S. Harischandra, H. Jin, V. Anantharam, A. Kanthasamy, A. G. Kanthasamy, alpha-Synuclein protects against manganese neurotoxic insult during the early stages of exposure in a dopaminergic cell model of Parkinson’s disease. Toxicol Sci 143, 454–468 (2015).

6. D. S. Harischandra et al., Manganese promotes the aggregation and prion-like cell-to-cell exosomal transmission of alpha-synuclein. Sci Signal 12 (2019).

7. S. Sarkar et al., Manganese activates NLRP3 inflammasome signaling and propagates exosomal release of ASC in microglial cells. Sci Signal 12 (2019).

8. D. S. Harischandra et al., Manganese-Induced Neurotoxicity: New Insights Into the Triad of Protein Misfolding, Mitochondrial Impairment, and Neuroinflammation. Front Neurosci 13, 654 (2019).

9. K. McFarthing et al., Parkinson’s Disease Drug Therapies in the Clinical Trial Pipeline: 2022 Update. J Parkinsons Dis 12, 1073–1082 (2022).

10. A. P. Ternes et al., Drosophila melanogaster - an embryonic model for studying behavioral and biochemical effects of manganese exposure. Excli J 13, 1239–1253 (2014).

11. S. Ghaisas et al., Chronic Manganese Exposure and the Enteric Nervous System: An in Vitro and Mouse in Vivo Study. Environ Health Perspect 129, 87005 (2021).

12. S. Sarkar et al., Manganese exposure induces neuroinflammation by impairing mitochondrial dynamics in astrocytes. Neurotoxicology 64, 204–218 (2018).

13. E. Belda et al., Impairment of gut microbial biotin metabolism and host biotin status in severe obesity: effect of biotin and prebiotic supplementation on improved metabolism. Gut 71, 2463–2480 (2022).

14. K. M. Lohr, B. Frost, C. Scherzer, M. B. Feany, Biotin rescues mitochondrial dysfunction and neurotoxicity in a tauopathy model. Proc Natl Acad Sci U S A 117, 33608–33618 (2020).

15. P. Ortega-Saenz et al., Selective accumulation of biotin in arterial chemoreceptors: requirement for carotid body exocytotic dopamine secretion. J Physiol 594, 7229–7248 (2016).

16. D. O. Kennedy, B Vitamins and the Brain: Mechanisms, Dose and Efficacy--A Review. Nutrients 8, 68 (2016).

17. S. Sarkar et al., Manganese exposure induces neuroinflammation by impairing mitochondrial dynamics in astrocytes. Neurotoxicology 10.1016/j.neuro.2017.05.009 (2017).

18. J. Kim et al., LRRK2 kinase plays a critical role in manganese-induced inflammation and apoptosis in microglia. PLoS One 14, e0210248 (2019).

19. E. Pajarillo et al., The role of microglial LRRK2 kinase in manganese-induced inflammatory neurotoxicity via NLRP3 inflammasome and RAB10-mediated autophagy dysfunction. J Biol Chem 299, 104879 (2023).

20. Q. J. Yu et al., Parkinson disease with constipation: clinical features and relevant factors. Sci Rep 8, 567 (2018).

21. H. Braak, U. Rub, W. P. Gai, K. Del Tredici, Idiopathic Parkinson’s disease: possible routes by which vulnerable neuronal types may be subject to neuroinvasion by an unknown pathogen. J Neural Transm (Vienna) 110, 517–536 (2003).

22. J. Horsager et al., Brain-first versus body-first Parkinson’s disease: a multimodal imaging case-control study. Brain 143, 3077–3088 (2020).

23. D. P. Breen, G. M. Halliday, A. E. Lang, Gut-brain axis and the spread of alpha-synuclein pathology: Vagal highway or dead end? Mov Disord 34, 307–316 (2019).

24. S. Kim et al., Transneuronal Propagation of Pathologic alpha-Synuclein from the Gut to the Brain Models Parkinson’s Disease. Neuron 103, 627–641 e627 (2019).

25. Z. D. Wallen et al., Metagenomics of Parkinson’s disease implicates the gut microbiome in multiple disease mechanisms. Nat Commun 13, 6958 (2022).

26. W. Yang et al., Current and projected future economic burden of Parkinson’s disease in the U.S. Npj Parkinsons Dis 6, 15 (2020).

27. M. Munzuroglu et al., Effects of biotin deficiency on short term memory: The role of glutamate, glutamic acid, dopamine and protein kinase A. Brain Res 1792, 148031 (2022).

28. X. Chen, C. Xie, L. Sun, J. Ding, H. Cai, Longitudinal Metabolomics Profiling of Parkinson’s Disease-Related alpha-Synuclein A53T Transgenic Mice. PLoS One 10, e0136612 (2015).

29. M. Langley et al., Mito-Apocynin Prevents Mitochondrial Dysfunction, Microglial Activation, Oxidative Damage, and Progressive Neurodegeneration in MitoPark Transgenic Mice. Antioxid Redox Signal 10.1089/ars.2016.6905 (2017).

30. V. Lawana et al., Involvement of c-Abl Kinase in Microglial Activation of NLRP3 Inflammasome and Impairment in Autolysosomal System. J Neuroimmune Pharmacol 10.1007/s11481-017-9746-5 (2017).

31. M. Neal et al., Prokineticin-2 promotes chemotaxis and alternative A2 reactivity of astrocytes. Glia 66, 2137–2157 (2018).

32. S. Sarkar et al., Mitochondrial impairment in microglia amplifies NLRP3 inflammasome proinflammatory signaling in cell culture and animal models of Parkinson’s disease. Npj Parkinsons Dis 3, 30 (2017).

33. S. Sarkar et al., Kv1.3 modulates neuroinflammation and neurodegeneration in Parkinson’s disease. J Clin Invest 130, 4195–4212 (2020).

34. R. Di Maio et al., alpha-Synuclein binds to TOM20 and inhibits mitochondrial protein import in Parkinson’s disease. Sci Transl Med 8, 342ra378 (2016).

35. D. G. Ordonez, M. K. Lee, M. B. Feany, alpha-synuclein Induces Mitochondrial Dysfunction through Spectrin and the Actin Cytoskeleton. Neuron 97, 108–124 e106 (2018).

36. S. Sarkar et al., Oligomerization of Lrrk controls actin severing and alpha-synuclein neurotoxicity in vivo. Mol Neurodegener 16, 33 (2021).

37. S. Brugman et al., A Comparative Review on Microbiota Manipulation: Lessons From Fish, Plants, Livestock, and Human Research. Front Nutr 5, 80 (2018).

38. S. Sarkar, M. Murphy, E. Dammer, A. Olsen, S. Rangaraju, E. Fraenkel, MB. Feany, Comparative proteomic analysis highlights metabolic dysfunction in α-synucleinopathy. Npj Parkinsons Dis In press (2020).

39. S. Sarkar, A. L. Olsen, K. Sygnecka, K. M. Lohr, M. B. Feany, alpha-synuclein impairs autophagosome maturation through abnormal actin stabilization. PLoS Genet 17, e1009359 (2021).

40. E. Hallacli et al., The Parkinson’s disease protein alpha-synuclein is a modulator of processing bodies and mRNA stability. Cell 185, 2035–2056 e2033 (2022).

41. A. O. Souvarish Sarkar, Katja Sygnecka, Kelly M. Lohr, Mel B Feany, α-synuclein impairs autophagosome maturation through abnormal actin stabilization. Plos Genetics Accepted in Press (2021).

42. S. Sarkar et al., Molecular Signatures of Neuroinflammation Induced by alphaSynuclein Aggregates in Microglial Cells. Front Immunol 11, 33 (2020).

43. Y. Lai et al., High-coverage metabolomics uncovers microbiota-driven biochemical landscape of interorgan transport and gut-brain communication in mice. Nat Commun 12, 6000 (2021).

44. H. Tsugawa et al., MS-DIAL: data-independent MS/MS deconvolution for comprehensive metabolome analysis. Nat Methods 12, 523–526 (2015).

45. H. Tsugawa et al., Hydrogen Rearrangement Rules: Computational MS/MS Fragmentation and Structure Elucidation Using MS-FINDER Software. Anal Chem 88, 7946–7958 (2016).

46. J. Ding et al., A metabolome atlas of the aging mouse brain. Nat Commun 12, 6021 (2021).

47. V. Neveu, G. Nicolas, A. Amara, R. M. Salek, A. Scalbert, The human microbial exposome: expanding the Exposome-Explorer database with gut microbial metabolites. Sci Rep 13, 1946 (2023).

48. E. L. Schymanski et al., Identifying small molecules via high resolution mass spectrometry: communicating confidence. Environ Sci Technol 48, 2097–2098 (2014).

49. Z. Pang et al., Using MetaboAnalyst 5.0 for LC-HRMS spectra processing, multi-omics integration and covariate adjustment of global metabolomics data. Nat Protoc 17, 1735–1761 (2022).

